# Proteomic Identification of Pig Xenoantigens for Clinical Xenotransplantation

**DOI:** 10.64898/2026.05.22.727249

**Authors:** Hongyi Liu, Trung Hoàng, Yingwei Hu, Yuanwei Xu, Zhenyu Sun, Brandon J. Peiffer, Yuanyu Huang, Zhaoli Sun, Hui Zhang

## Abstract

Xenotransplantation using genetically engineered pig organs offers a promising solution to the shortage of donor organs for life-saving transplants. However, human preformed antibodies against unknown pig xenoantigens remain a significant barrier to successful xenotransplantation. Current methods for characterizing these antibodies or xenoantigens are limited to cellular-level crossmatch assays. In this study, we developed a novel approach to identify pig xenoantigens, including peptide and glycopeptide epitopes that react with human preformed antibodies. First, human preformed antibodies against xenoantigens were enriched from plasma using immobilized pig kidney proteins. The enriched antibodies were then immobilized and used to isolate pig kidney proteins, peptides, and intact glycopeptides, followed by liquid chromatography-tandem mass spectrometry analysis. This dual-level approach identified 221 peptides corresponding to 153 proteins, with a significant enrichment of plasma membrane and extracellular proteins. Notably, 11 peptides were unique to pig sequences, suggesting their potential role in driving xenogeneic immune responses. Glycoproteomic analysis identified 122 intact glycopeptides, predominantly complex/hybrid glycoforms and Neu5Gc-containing glycans. Our method effectively identifies peptides and intact glycopeptides reactive to human preformed antibodies, providing critical insights for discovering xenoantigens. These findings could guide genetic engineering strategies and enhance recipient candidate screening for xenotransplantation, ultimately increasing the feasibility and success of xenogeneic organ transplantation.

## INTRODUCTION

Xenotransplantation, the transplantation of organs from non-human species, particularly pigs, has been proposed as a potential solution to the shortage of human donor organs^1–3^. However, a significant challenge to the clinical success of xenotransplantation is the presence of natural (preformed) human antibodies that recognize pig xenoantigens, leading to antibody-mediated rejection of pig organ grafts^4,5^. Extensive research has identified three primary glycan xenoantigens: Galactose-α1,3-galactose (Gal), N-glycolylneuraminic acid (Neu5Gc), and Sda, which are recognized by human anti-pig antibodies^6–8^. To overcome this, genetically engineered ‘triple-knockout’ (TKO) pigs that lack these antigens were developed^9^. Using TKO pigs in combination with advanced co-stimulation blockade-based immunosuppression (IS), significant graft survival has been achieved in nonhuman primate (NHP) models across multiple organ systems^10^. Encouraged by these results, genetically engineered pig hearts and kidneys were transplanted into human recipients under compassionate use circumstance^11,12^. However, these clinical cases resulted in limited success, with either patient death or xenograft removal within months post-transplantation^13^.

A two-month study involving the transplantation of genetically engineered pig kidneys into brain-dead human recipients revealed no hyperacute rejection^14^. However, detailed phenotyping of the xenografts indicated early signs of antibody-mediated rejection^15^, including microvascular inflammation, immune deposits, endothelial cell activation, and positive xenoreactive crossmatches. These observations suggest the existence of additional xenoantigens beyond the known triple glycan antigens that are recognized by human antibodies. This hypothesis is supported by findings showing that over 60% of human sera contain antibodies binding to TKO pig cells, with 40% of human sera IgM cross-reacting at low levels^16^. Additionally, swine leukocyte antigens (SLA) class I and II can be recognized by cross-reactive human leukocyte antigen (HLA) antibodies^17^.

Current methods for detecting anti-pig antibodies predominantly utilize flow cytometry crossmatch assays that measure antibody binding to pig red blood cells (RBCs) or peripheral blood mononuclear cells (PBMCs)^18–20^. However, these methods have limitations. For example, antibody binding to pRBCs reflects humoral reactivity to carbohydrate xenoantigens but does not provide insights into immunologic sensitivity to pig protein antigens such as SLAs, which are absent on pRBCs. Conversely, pPBMCs are more suitable for evaluating overall humoral responses to both carbohydrate and protein antigens but fail to represent all glycoproteins and proteins expressed across different pig organs. Organ-specific glycoproteins possess unique glycan structures that can vary considerably between tissues^21^. Consequently, the current antibody detection methods, while crucial for patient selection in clinical trials, are limited by their reliance on cell-based crossmatch assays and the absence of commercially available solid-phase assays^4,22,23^.

Recent advancements in mass spectrometry have significantly enhanced proteomics technologies^24,25^. Notably, Data-Independent Acquisition (DIA) technology enables high-sensitivity and reproducible identification of low-abundance proteins in complex samples^26,27^. Additionally, improvements in enrichment methods and search engines have advanced the identification of glycosylated proteins^28,29^, which play a pivotal role in immune responses to xenoantigens. Traditional antigen discovery methods, such as immunoprecipitation, rely on antibody-antigen interactions^30,31^, which can be confounded by non-specific protein-protein interactions, impacting the reliability of antigen identification^32,33^. Unlike full-length proteins, antibodies can bind to specific short sequences on proteins known as epitopes. Peptides, which form these epitopes, exhibit minimal non-specific interactions^34,35^, making peptide-level enrichment a promising approach for enhancing the specificity of xenoantigen identification.

This study aimed to identify pig xenoantigens that react with human preformed antibodies using an innovative proteomic approach. We developed a method that integrates the enrichment of human preformed antibodies binding to pig proteins and the isolation of peptide epitopes recognized by these antibodies. This was followed by liquid chromatography-tandem mass spectrometry (LC-MS/MS) analysis, providing a robust and reliable strategy for identifying pig xenoantigens. Our findings revealed not only the cellular localization of pig kidney-derived xenoantigens but also highlighted the immunogenic potential of glycoproteins and peptides that differ from human sequences. Furthermore, we identified several intact glycopeptides with specific glycosylation patterns, which could be key targets for mitigating xenogeneic immune responses. These results underscore the importance of future studies focusing on the immunogenic properties of these glycopeptide and peptide epitopes, paving the way for novel strategies to enhance xenotransplantation outcomes. This study contributes to the ongoing effort to identify new xenoantigen targets, optimize the genetic engineering of donor pigs, improve xenotransplant recipient selection, and develop innovative therapeutic interventions to enhance graft survival and clinical success in xenotransplantation.

## EXPERIMENTAL SECTION

### Pig kidney tissue lysis

The lysis buffer was prepared as 1 mM ethylenediaminetetraacetic acid (EDTA), 20 uM O-(2-Acetamido-2-deoxy-D-glucopyranosylidenamino) N-phenylcarbamate (PUGNAc) (Sigma, A7229), and 1X radioimmunoprecipitation assay buffer (RIPA) buffer diluted from 10X RIPA buffer (Merck, 20-188). This mixture was kept on ice. Next, 200 ul of the chilled lysis buffer was added to 50 mg of cryopulverized pig kidney tissue recovered from Yorkshire swine obtained from a commercial vendor. The tissue and lysis buffer mix were vortexed for 20 seconds on the highest setting. The tissues were then lysed at 4 °C with the chilled lysis buffer for 15 minutes. This process was repeated two more times for a total of three cycles. The mixture was then transferred to a 2 ml Eppendorf tube labeled as an insoluble pellet and centrifuged at 4 °C (20,000 g) for 10 minutes to remove cell debris. The supernatant was then transferred to a 2 ml conical tube, measuring the amount of supernatant while keeping the insoluble pellet. The protein concentration was estimated with a bicinchoninic acid assay (BCA) assay using the supernatant. Samples, a water blank, and a lysis buffer sample were diluted 1:20 and run in triplicate. The samples were incubated at 37 °C for 30 minutes in a Thermomixer and then the plate was read at 562 nm.

### Pig kidney protein linked beads generation

We converted the pig kidney protein buffer into phosphate-buffered saline (PBS) using ultrafiltration (100 ul diluted to 400 ul, 10 times) to avoid the presence of tris or other buffers containing primary amines. Then, we transferred the AminoLink Plus coupling resins (Thermo Scientific, 20501) to a new tube, with 1 ml of resins used for each 10 mg of proteins. The resins were washed with PBS, vortexing, and centrifuging, with the supernatant discarded. This process was repeated three times. The pig kidney proteins were then transferred to the washed resin and incubated overnight at 4 ℃ with gentle rotation, creating pig kidney protein beads (PKP beads). Following incubation, the sample was centrifuged to collect the flow-through, which was used to determine the coupling efficiency by comparing protein concentrations with the starting sample. The PKP beads were then washed with PBS twice. Next, in a fume hood, NaBH_3_CN was added to a final concentration of 50 mM in PBS and incubated for 4 hours at room temperature. After centrifugation, the PKP beads were washed with 1 M Tris-HCl and centrifuged. This step was repeated once. The unused aldehyde groups on the PKP beads were blocked by adding NaBH_3_CN to Tris-HCl and incubating for 30 minutes with gentle rotation at room temperature. After centrifugation, the flow-through was discarded, and the reactants and non-coupled protein were washed away with 1 M NaCl. The PKP beads were then equilibrated for storage by adding PBS and washing twice.

### Preformed human antibody purification

The PKP beads were transferred to a 5 ml tube with empty AminoLink Plus coupling resins as negative controls, and 5 ml pooled human plasma (Innovative Research, IPLAWBK2E100ML) was added to each tube. The samples were incubated overnight at 4 ℃. After centrifugation, the flowthroughs were saved and the PKP beads were transferred to a column. The resins were washed with PBS and centrifuged 4 times. Human pre-formed antibodies reactive to pig xenoantigens were eluted with 0.1% TFA, incubated for 10 minutes, and then centrifuged. This step was repeated three times. The eluted xenoantibodies were neutralized with PBS. After use, the PKP beads were equilibrated with 0.1 M sodium phosphate pH 7.4 to prevent damage to the immobilized protein by the low pH elution buffer.

### Human preformed antibody conjugation to beads

The 10 ul gel/isopropanol slurry of the Affi-Gel Hz Hydrazide Gel (Bio-Rad, 156-0017) was transferred to a clear 0.5 ml Eppendorf tube and allowed to settle. The isopropanol supernatant was removed, and the gel was mixed with 200 ul of diluted coupling buffer, pH 5.5. This was repeated once more. Unused gel was stored at 4 °C in isopropanol. Sodium periodate was dissolved in water and added to the purified 30 ug human pre-formed antibody samples with a final concentration at 10 mM. The mixture was rotated end-over-end for 1 hour at room temperature, then immediately desalted by ultracentrifuge. This light-sensitive reaction was performed in the dark. The oxidized, desalted preformed antibody samples were added to the gel in the 0.5 ml Eppendorf tube and rotated end-over-end for overnight at room temperature. After the coupling reaction was complete, the supernatants were collected for efficiency determination. The xenoantigen reactive antibody linked beads (Ig beads) was washed with PBS and centrifuged. In a fume hood, NaBH_3_CN was added to a final concentration of 50 mM in PBS and incubated for 4 hours at room temperature. The Ig-beads were then washed with a buffer containing 0.5 M NaCl PBS twice, followed by PBS twice. The Ig beads were aliquoted to 2 portions for protein or peptide-level enrichment and stored at 4 °C. Freezing was avoided to maintain the integrity of the beads.

### Protein-level enrichment

The first portion of Ig beads were first washed with 200 ul of PBS twice. After this, 500 ug of pig proteins are aliquoted and diluted to 100 ul with PBS, then added to the 5 ul Ig beads. The mixture was incubated overnight with moderate agitation at 4 ℃. After the incubation period, the Ig beads were washed twice with 200 ul of 0.5 M NaCl in PBS and then washed twice with PBS. Following these washings, 20 ul of 1% Formic Acid (FA) was added to the beads, and the mixture was incubated for 10 minutes with moderate agitation, the eluate was freeze dried and keep frozen at −80 ℃ until use.

### In solution digestion of enriched pig proteins

To digest the eluted proteins in solution, proteins were solved in 2 ul lysis buffer containing 0.1% n-Dodecyl-B-D-maltoside (DDM), 5 mM tris(2-carboxyethyl)phosphine (TCEP), and 20 mL iodoacetamide (IAA) in 100 mM tetraethylammonium bromide (TEAB). The proteins were reduced and alkylated with for 1 hour in the dark at 80 ℃. Next, 1 ug lysC and 1 ug trypsin were added and incubated overnight at 37 °C. The lysC and trypsin mix stock used was 1 ug/ul. Finally, 50% FA was added to acidify the digest with 1% FA in the final solution, bringing the pH to 2.5. The sample was then diluted up to 100 ul with 0.1% FA. The sample was centrifuged at 12,000 g, for 15 minutes and the supernatant was recovered for the subsequent analyses.

### Peptide-level enrichment

The second portion of Ig beads were first washed with 200 ul of PBS twice. After this, 200 ng of pig peptides were aliquoted and diluted to 10 ul with PBS, then added to the Ig beads. The mixture was incubated overnight with moderate agitation at 4 ℃. After the incubation period, the beads were washed twice with 200 ul of PBS. Following these washings, 20 ul of 1% FA was added to the beads, and the mixture was incubated for 10 minutes with moderate agitation, the eluate was lyophilized and frozen at −80℃. The samples were then resuspended in 30 ul of 0.1% FA, and 50 ng of the sample was used for LC-MS analysis.

### Preparing peptides for LC-MS/MS analysis

The procedure started with conditioning an Evotip with 100 ul acetonitrile (MeCN), followed by soaking in isopropyl alcohol (IPA) for 10 seconds. The cartridge was then equilibrated with 100 ul of 0.1% FA twice. Following this, the prepared sample was loaded onto the Evotip. The Evotip was then washed and desalted with 100 ul of 0.1% FA. After that, the sample was eluted from the cartridge with 50 ul of 50% MeCN /0.1% FA. The eluate was then frozen either in liquid nitrogen or at −80℃ and lyophilized. The samples were resuspended in 0.1% FA and 50 ng samples were injected into Ascend for LC-MS analysis and 200 ng samples used for peptide-level enrichment.

### EvoSep for global and glycosylation proteomic analysis

The pig kidney tissue LC-MS/MS data were acquired via EvoSep coupled with timsTOF HT (Bruker) in data-independent acquisition mode. The methods for acquiring global proteomics are as follows. Global peptides were loaded on Evotip and separated on PepSep C18 column of 15 cm x 150 µm, 1.5 µm (Bruker) in Bruker column toaster (50 °C) at a 30 samples per day (SPD) gradient. The global proteomic data was acquired using the DIA-PASEF mode on timsTOF-HT with settings as follows: MS1 scan range of 100-1700 m/z; MS2 scan range of 338-1338 m/z, mass width 25.0 Da without mass overlap, 1 mobility window, 1/K0 range of 0.70-1.45 V.s/cm2, ramp time 85.0 ms.

The pig kidney xenoantigen enriched by protein and peptide-level global LC-MS/MS data were acquired via EvoSep coupled with Ascend (Thermo Scientific) in DIA acquisition mode. The global proteomic data was acquired using the DIA mode on Ascend with settings as follows: MS1 scan range of 380-985 m/z; MS2 scan range of 145-1450 m/z, isolation window 18.0 m/z with mass overlap 1 m/z. Stepping HCD energy 22, 27, 32 was utilized here.

The intact glycopeptides of pig kidney xenoantigen enriched by peptide-level enrichment were analyzed by data dependent acquisition (DDA) on Ascend (Thermo Scientific) combined with EvoSep. The ion source of the mass spectrometer was set up with 1.8 kV of electrospray voltage and 300 °C of ion transfer tube temperature. The precursor ion scan was acquired with 120 K resolution at 200 m/z for from 350 to 2,000 m/z range and AGC value was set as 5 × 10^5^. Precursor ions isolated from 0.7 m/z width were fragmented by stepping HCD energy 25, 35, 45 and fragment ions were acquired with 30 K resolution with 3 × 10^5^ of AGC value for 64 ms of injection time for a total duty cycle (2 s). The peptide charge state screening was enabled to include 2 to 8 ions with a dynamic exclusion time of 45 s to discriminate against previously analyzed ions between +/− 10 ppm.

### Data Processing

For DIA proteomic data searching, spectral libraries were generated in Spectronaut® 18.4 (Biognosys AG) by combination of search archives from pig kidney tissue samples or pig xenoantigen samples. All the raw files were searched against the pig protein database downloaded from the UniProt/Swiss-Prot (version, 2024-3) database, which was appended with an equal number of decoy sequences in the Spectronaut. The search setting is as follows. The mass tolerance of MS and MS/MS was set as dynamic with a correction factor of one. Precursors were filtered by a Q value cutoff of 0.01 which corresponds to a false discovery rate (FDR) of 1%. Carbamidomethyl (C) was set as a fixed modification. Acetyl (Protein N-term) and Oxidation (M) were set as variable modifications. The quantity of a peptide was a sum of the quantity of its top 3 precursors, whereas the quantity for a precursor was calculated by summing the area of its top 3 fragment ions at the MS/MS level. The data was normalized by being divided by the median intensity value of each sample.

For DDA glycoproteomic data searching, prior to database searching, ProteoWizard 3.0 was used to convert .RAW files to .mzML format with the “peakPicking” option enabled. The data were analyzed using GPQuest (version 3.0) on a local computing system (64 vCPUs, 1 TB RAM)^28^. A mammalian N-glycan database containing 309 glycans, pre-processed to remove sodium adducts, and a customized pig kidney protein database with 7,800 sequences from UniProt were used for glycoproteomic identification. Search parameters included a precursor mass tolerance of 10 ppm and a fragment mass tolerance of 20 ppm. Trypsin was specified as the digestion enzyme, allowing up to two missed cleavages, with peptide sequences restricted to lengths between 7 and 50 amino acids and precursor charge states ranging from +2 to +8. For peptide fragment ions, b- and y-type ions were considered, with a minimum of 6 matched peaks required for identification. Carbamidomethylation of cysteine (+57.0215 Da) was set as a fixed modification, while methionine oxidation (+15.9949 Da) was considered a variable modification. The results were filtered using a 1% PSM FDR threshold. Unless otherwise stated, data are presented as mean ± standard error of the mean. Data are derived from experiments repeated at least three times unless stated otherwise. GraphPad Prism 8 and R 4.2.2 were used for statistical analysis.

## RESULT

### Workflow of protein and peptide-level enrichment by using the immobilized antibodies from human plasma

We developed a proteomic method to identify and quantify xenoantigens that were recognized by human pre-formed antibodies. This approach involved pre-antibody and pig xenoantigen enrichment strategies, followed by LC-MS/MS analysis. Initially, antibodies specific to pig kidney proteins were immunoprecipitated. The immunoprecipitation process involved isolating pre-formed antibodies from human plasma using immobilized pig proteins as the target (Figure 1). To determine the optimal ratio of pig protein linked beads to human pooled plasma for antibody enrichment, we conducted a series of tests varying both the volume of pig kidney protein (PKP) beads and the volume of plasma (Figure S1 and Supplementary table S1). Our findings indicated that the most effective ratio of PKP beads to plasma was approximately 1:10. This ratio optimized the beads’ binding capacity. The antibodies were then covalently bound to hydrazide beads. In the next phase, pig kidneys were lysed and the proteins incubated with the antibody-bound hydrazide beads, allowing pig kidney proteins to be captured (protein-level enrichment) (Figure 1). Following the protein-level enrichment, the captured proteins were digested into peptides. These peptides were then incubated with the antibody-bound hydrazide beads, allowing specific peptide epitopes to be captured (peptide-level enrichment). The enriched peptides were then subjected to LC-MS/MS analysis. Overall, this workflow combining immunoprecipitation, covalent bead attachment, dual-level enrichment, and LC-MS/MS analysis ensured specific identification of pig xenoantigens bound to human pre-antibodies.

**Figure 1.**
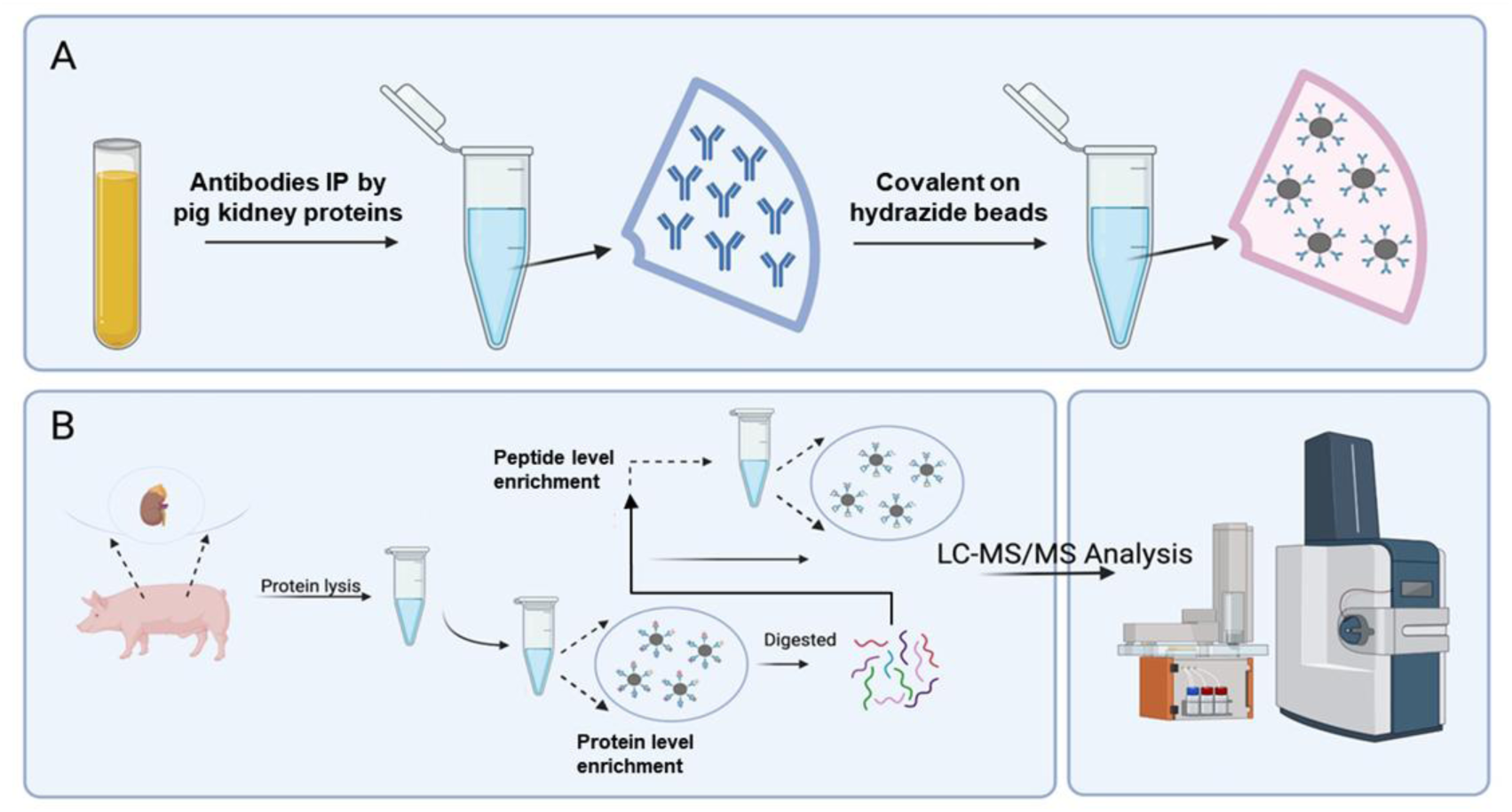
Workflow of protein and peptide-level enrichment by using the immobilized antibodies from human plasma. A. Antibodies were immunoprecipitated (IP) by pig kidney proteins and then covalently bound to hydrazide beads. B. Proteins from pig tissues were lysed, captured by the antibody-bound beads (protein-level enrichment) and then digested to peptide and go through the peptide-level enrichment. These captured proteins or peptides were then analyzed using LC-MS/MS to identify the xenoantigens.

### The reproducibility and quantification of pig xenoantigens bound to human preformed antibodies

Assessing the repeatability and quantification of xenoantigens is a critical step in ensuring the reliability of this approach to xenotransplantation research. The correlation analysis of xenoantigen enrichment provided insights into the enrichment patterns and the specificity of antibody interactions with pig kidney xenoantigens. We compared the correlation of samples enriched without xenoantigen-specific antibodies (E group, E1-E3) and those enriched with antibodies bound to pig kidney proteins (PK group, PK1-PK3). The E group consisted of beads without xenoantigen-specific antibodies, which were pulled down from human plasma by empty beads without conjugated pig kidney proteins, while the PK group comprised beads with antibodies pulled down from human plasma by pig kidney proteins (Figure 1A). For protein-level enrichment, the results indicated strong correlations within each group, as evidenced by the uniform high correlation coefficients among the E1, E2, and E3 samples, as well as among the PK1, PK2, and PK3 samples (Figure 2A), indicating consistent enrichment patterns within each group (Figure 2A). Importantly, the correlations between the E group and the PK group were lower in the peptide-level analysis compared to the protein-level analysis (Figure 2A and B). This showed that the antibodies in the PK group specifically target distinct peptides derived from pig kidney proteins, resulting in unique enrichment patterns that differ from those observed without xenoantigen-specific antibodies.

**Figure 2.**
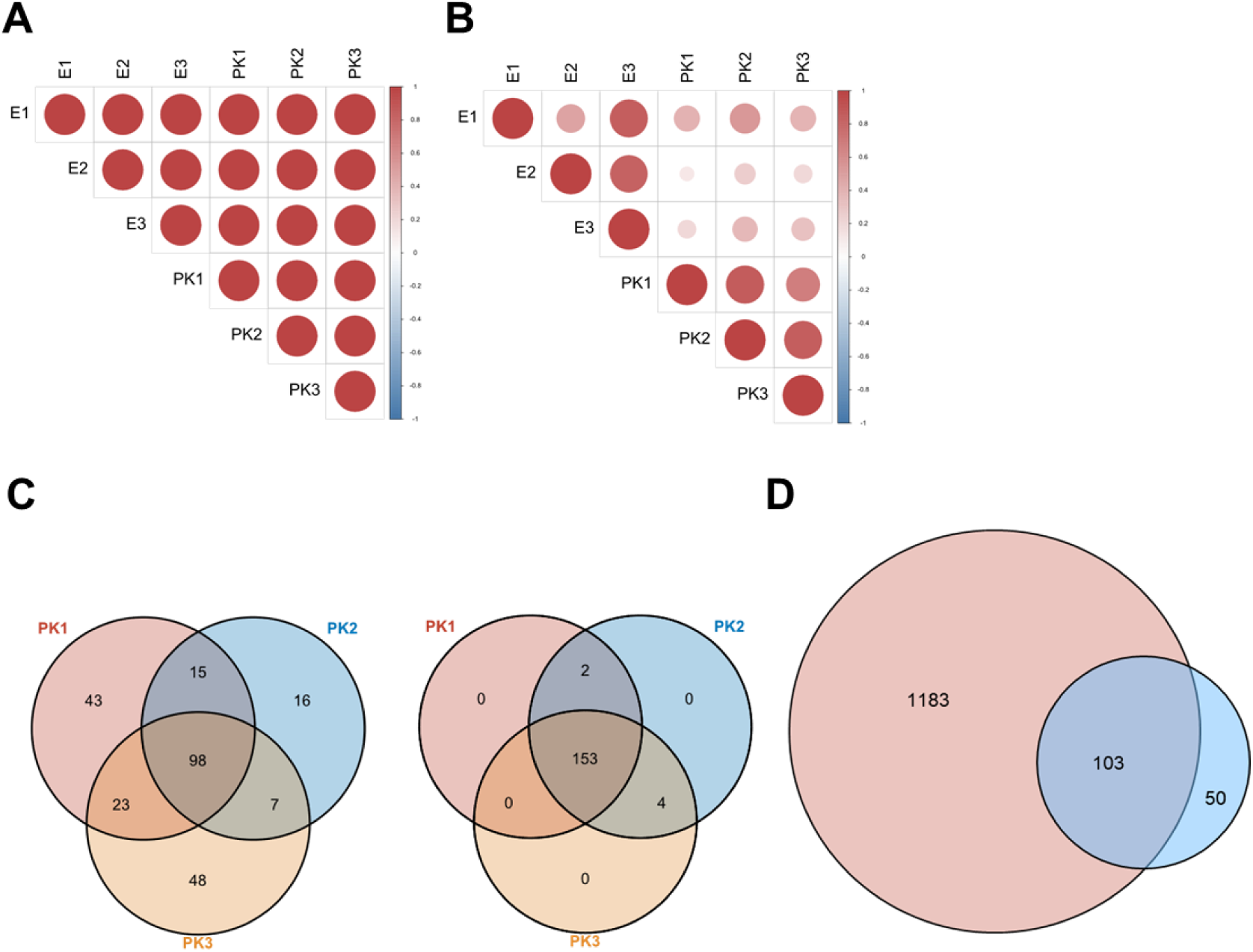
The reproducibility and quantification of pig xenoantigens bound to human preformed antibodies. A and B. The correlation of protein (A) and peptide (B) -level enrichment results. The E group were beads with non-specific antibodies from human plasma. The PK group were beads with antibodies IP by pig kidney proteins from human plasma. C. Venn diagram illustrating the overlap of proteins enriched across three replicate samples (PK1, PK2, PK3) at the protein-level enrichment (left) or peptide-level enrichment (right). D. Venn diagram depicting the overlap between proteins identified at the protein-level enrichment (red, 1,286 proteins) and proteins enriched at the peptide-level enrichment (blue, 153 proteins).

We further performed quantification analysis of both protein and peptide-levels. The results showed significant non-specific binding in the protein-level enrichment, as evidenced by the overlap between the fold changes of the PK group (beads with specific antibodies to pig kidney proteins) and fold changes of the E group (beads without specific antibodies) distributions (Figure S2A). This overlap suggested that protein-protein interactions contributed to high non-specific binding, potentially confounding the identification of real xenoantigens. In contrast, the peptide-level enrichment displayed a markedly different pattern (Figure S2B). The intersections of the fitting curves were distinct, indicating that the peptide-level enrichment had high specificity. There were fewer overlaps, suggesting that non-specific binding was almost negligible. Most peptides in the PK group were upregulated compared to the E group. These findings underscored the advantages of peptide-level enrichment over protein-level enrichment in terms of specificity.

To assess the reproducibility and specificity of our enrichment strategies, we quantified protein and peptide-level enrichments and presented the results using Venn diagrams. The result showed the overlap of proteins enriched across three replicate samples (PK1, PK2, PK3) (Figure 2C). A total of 98 proteins were commonly enriched in all three replicates. However, there were also several proteins unique to each replicate, PK1 had 43 unique proteins, PK2 had 16, and PK3 had 48, suggesting variability that could be attributed to non-specific protein-protein interactions. For the peptide-level enrichment, a total of 251 peptides from 153 proteins were consistently enriched in all three replicates, highlighting the high reproducibility of the peptide-level enrichment method (Figure 2C and Supplementary table S2). Unlike protein-level enrichment, the peptide-level enrichment showed minimal variability, with few peptides unique to individual replicates. This indicated that peptide-level enrichment was more specific and consistent compared to protein-level enrichment. The overlap of 103 proteins between protein-level identified (1,286 proteins) and peptide-level enriched (153 proteins) emphasized the broad coverage of protein-level enrichment (Figure 2D). The clear distinction between protein and peptide-level identifications suggested that peptide-level enrichment provided a more specific and reliable approach for identifying xenoantigens, minimizing the confounding effects of non-specific protein interactions.

### Intersections of protein and peptide-level enriched xenoantigens across cellular components

To elucidate the cellular localization and potential immunogenicity of enriched proteins derived from pig kidney, we conducted an intersection analysis across cellular components and performed sequence alignment comparisons between pig and human protein sequences. The intersection analysis revealed that in the protein-level enrichment result, 34 proteins were localized to the plasma membrane, indicating significant enrichment of membrane-associated proteins (Figure 3A). At the peptide level, 29 proteins were identified in the plasma membrane, demonstrating focused enrichment of membrane-bound peptides (Figure 3B). Both protein and peptide-level enrichments successfully highlighted antigens from the plasma membrane and extracellular regions, critical for identifying potential xenoantigens. For the protein-level enriched proteins, 32 of them are glycoproteins, and for the peptide-level enriched proteins 7 of them are glycoproteins according to the UniProt database (Supplementary table S3).

**Figure 3.**
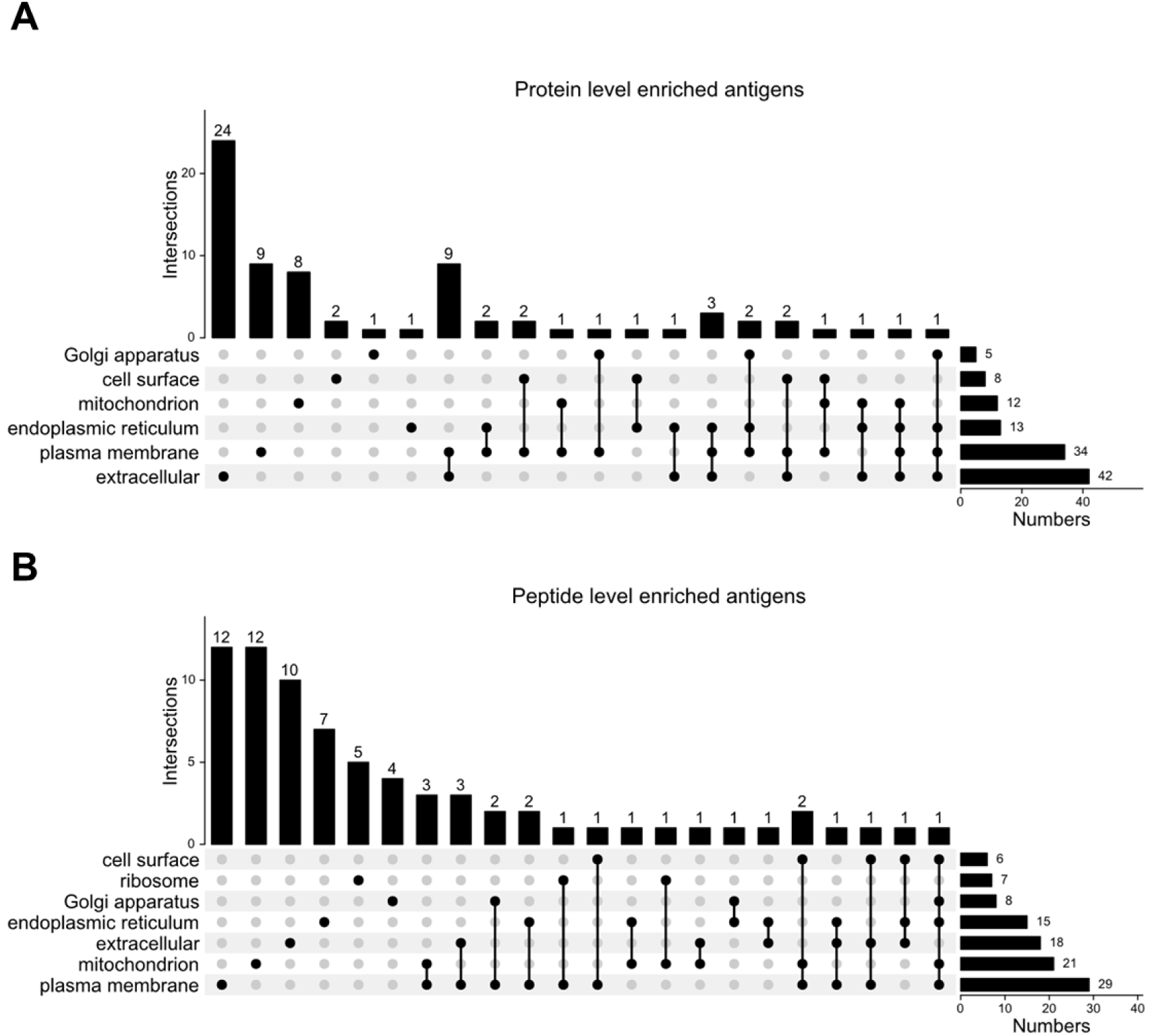
Intersections of protein and peptide-level enriched xenoantigens across cellular components. A. Intersection analysis of protein-level enriched xenoantigen candidates across various cellular components. The bar plot showed the number of proteins found at each intersection of cellular components. B. Intersection analysis of peptide-level enriched xenoantigen candidates across various cellular components. The bar plot showed the number of proteins found at each intersection of cellular components.

To assess immunogenicity, sequence alignment analysis compares identified peptide sequences between pig and human. The results indicated that 208 sequences were identical to those in human proteins (Supplementary table S3), suggesting a lower likelihood of triggering an immune response. In contrast, 11 pig peptides differed from human sequences and these mismatches may drive xenogeneic immune reactions (Supplementary table S3). These findings highlight the importance of further investigating these peptide sequences to understand their roles in immune response mechanisms.

### Glycopeptide analysis of the peptide-level enrichment

To identify intact glycopeptides in the peptide-level enrichment, we first performed label-free DIA proteomics on pig kidney peptides. In total, we identified 7,439 protein groups in pig kidney (Supplementary table S4). These proteins were then used as the FASTA database for intact glycopeptide identification. Next, we performed label-free DDA intact glycoproteomic analysis on peptide-level enriched peptides. In three replicates of the PK group with specific enrichment by xenoantibodies, we identified 122 intact glycopeptides, 33 of which were consistently identified in all three replicates. Among these, 31 intact glycopeptides belonged to haptoglobin (Supplementary table S4). By contrast, in the E group with non-specific enrichment, no intact glycopeptides were identified in all three replicates.

The distribution of glycoforms for the 122 intact glycopeptides in the PK group, highlighting the predominance of complex/hybrid glycoforms, with some contributions of high mannose glycoforms (Figure 4A). Importantly, the frequency of sialic acid types showing not only the Neu5Ac as the sialic acid, but also with Neu5Gc present in 17 intact glycopeptides (Figure 4B). These findings highlight the specificity of xenoantibody-based enrichment in capturing intact glycopeptides, particularly those containing Neu5Gc glycans.

**Figure 4.**
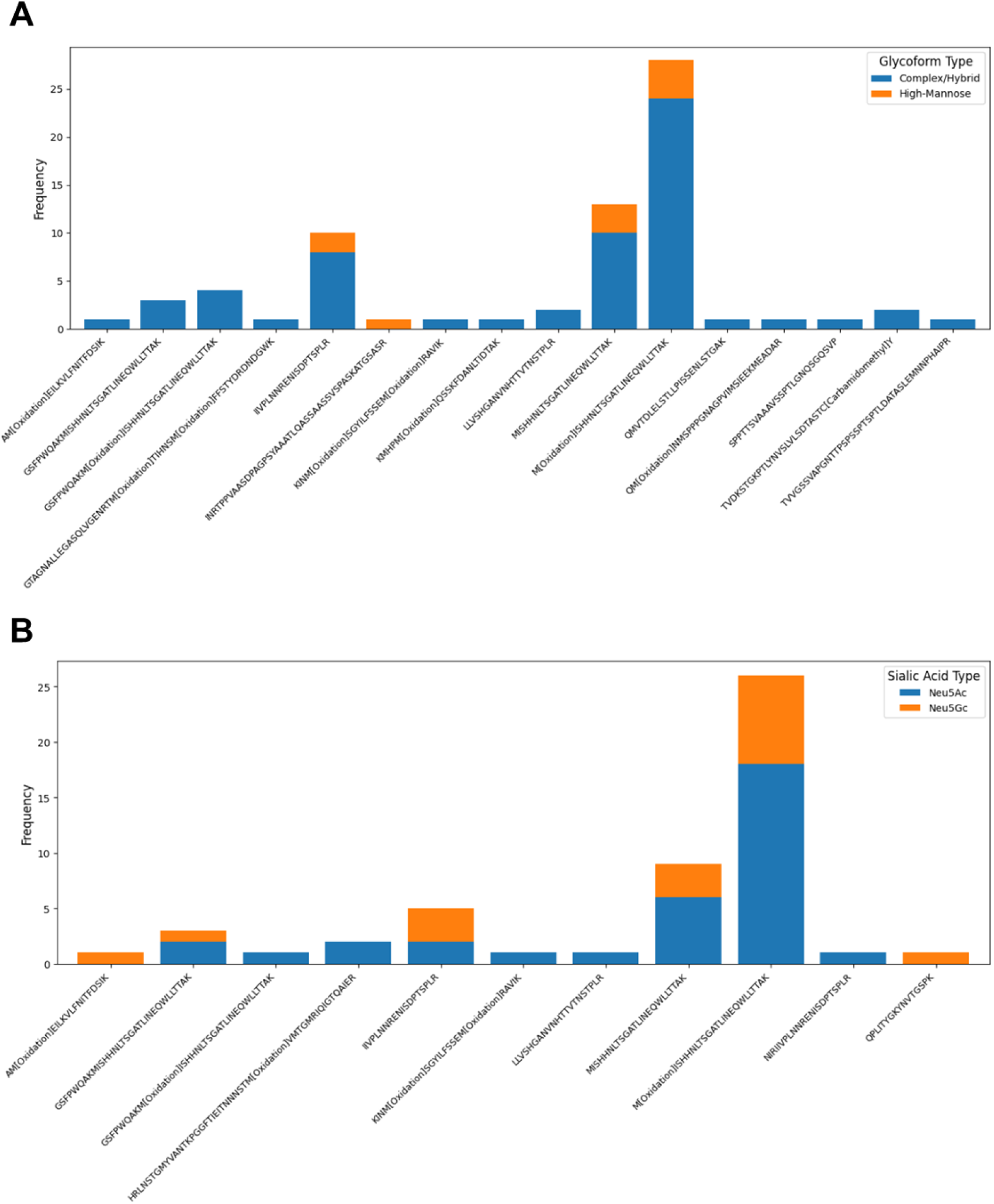
Glycopeptide analysis of the peptide-level enrichment. A. The bar graph illustrated the frequency distribution of intact glycopeptides categorized by glycoforms, including complex/hybrid (blue) and high mannose (orange). B. The bar graph illustrated the frequency distribution of intact glycopeptides based on sialic acid types, Neu5Ac (blue) and Neu5Gc (orange).s

## DISCUSSION

In this study, we developed a novel dual-level proteomic approach in which enriched human preformed antibodies from human plasma using immobilized pig proteins were used to isolate pig kidney proteins and peptide epitopes. This was followed by liquid chromatography-tandem mass spectrometry analysis, aiming to identify pig xenoantigens that react with human preformed antibodies. Using this approach, we identified 221 peptides corresponding to 153 proteins, with a significant enrichment of plasma membrane and extracellular proteins. Notably, 11 peptides were unique to pig sequences, suggesting their potential role in driving xenogeneic immune responses. Glycoproteomic analysis identified 122 intact glycopeptides, predominantly complex/hybrid glycoforms and Neu5Gc-containing glycans. Our findings provide new insights into identifying xenoantigens that react with human preformed antibodies.

The dual-level enrichment approach maximized the likelihood of capturing xenoantigens by leveraging concentration differences between antibodies and antigens. Antibody-antigen interactions are concentration-dependent and follow the principles of chemical equilibrium and affinity binding. Using highly concentrated pig proteins immobilized on AminoLink beads to enrich antibodies from human serum allowed for the capture of human preformed antibodies against pig antigens present at lower levels. Meanwhile, immobilizing these specific antibodies onto hydrazide beads highly enriched antibody presentation by immobilization of human preformed antibodies through carbohydrates presented at the constant region of antibodies and leaving variable region of preformed antibodies available for xenoantigen binding, thereby increasing the sensitivity of xenoantigen capture^36–38^.

The dual-level enrichment approach ensured the specificity of xenoantigen and human preformed antibody interactions. Correlation analysis between samples enriched without specific antibodies (E group) and those with antibodies specific to pig kidney proteins (PK group) demonstrated the robustness of our enrichment strategies. Strong within-group correlations observed in protein and peptide-level enrichment (E1-E3 and PK1-PK3) underscore the consistency of antibody interactions with pig kidney xenoantigens. The lower correlations between the E and PK groups at the peptide level highlight the specificity of the antibodies in the PK group, which target distinct peptides derived from pig kidney proteins. Non-specific binding observed in protein-level enrichment, evidenced by the overlap in fold changes between the PK and E groups, suggests that protein-protein interactions contribute substantially to non-specific binding at the protein level. This overlap could potentially confound the identification of true xenoantigens. Conversely, the distinct separation in fold changes at the peptide level indicated high specificity and minimal non-specific binding. Most peptides in the PK group were upregulated compared to the E group, underscoring the effectiveness of peptide-level enrichment in isolating specific antibodies against pig kidney proteins. Venn diagrams quantifying the reproducibility of our enrichment strategies showed a total of 98 proteins commonly enriched across all three replicates in protein-level enrichment, with variability attributed to non-specific protein-protein interactions. In contrast, peptide-level enrichment demonstrated high reproducibility, with 153 proteins consistently enriched across all replicates and minimal variability. The substantial overlap of 103 entities between protein-level identified (1,286 proteins) and peptide-level enriched (153 proteins) emphasized the broad coverage of protein-level enrichment while highlighting the specificity of peptide-level enrichment. Additionally, glycoproteomic analysis on peptide-level enriched peptides led to the identification of 122 intact glycopeptides, while no intact glycopeptides were consistently identified across three replicates in the E group with non-specific enrichment, indicating the specificity of our enrichment strategy.

The well-known xenoantigens in pig tissues that trigger antibody-mediated rejection in humans and other primates are glycan structures present on glycoproteins and glycolipids, including α-Gal (Galα1-3Galβ1-4GlcNAc-R), N-Glycolylneuraminic Acid (Neu5Gc), and SDa-like Glycans (GalNAcα1-4Gal-R)^16^. The identification of 17 intact glycopeptides containing Neu5Gc glycans was particularly noteworthy. Neu5Gc is a sialic acid present in pigs but absent in humans due to a genetic mutation in the *CMAH* gene^39–41^. Anti-Neu5Gc antibodies in humans contribute to chronic rejection, inflammation, and hyperacute rejection in xenotransplantation^42^. In addition, our findings also suggest the presence of classic glycoforms known to function as xenoantigens, such as the α-Gal epitope and the carbohydrate antigen Sda (a terminal GalNAc residue β4-linked to an α3-sialylated galactose) in identified intact glycopeptides. Although the identical molecular weights of glycans in humans and pigs prevent definitive conclusions, the presence of glycan compositions capable of generating these two well-known xenoantigen glycoforms (knocked out in genetically engineered TKO pigs) in our identified results provides strong evidence for the reliability of our method.

The intersection analysis of xenoantigens in protein and peptide-level enrichments across various cellular components, along with sequence alignment comparisons between pig and human protein sequences, provided insights into the cellular localization and potential immunogenicity of enriched proteins derived from pig kidney. By identifying the specific cellular compartments where these proteins were localized, we can better understand their roles and potential as targets in xenotransplantation. Our findings revealed that a substantial number of proteins identified through protein-level enrichment were associated with the plasma membrane (34 proteins) and extracellular region (42 proteins). Similarly, peptide-level enrichment also identified proteins localized to the plasma membrane (29 proteins) and extracellular region (21 proteins). These results underscore the ability of our enrichment strategies to effectively target membrane-bound and extracellular proteins, which are crucial for xenoantigen identification due to their accessibility and potential immunogenic properties. Sequence alignment analysis between pig and human proteins revealed that 208 peptide sequences were identical, suggesting a lower likelihood of these sequences triggering an immune response. However, 11 pig peptides exhibited sequence differences from human proteins, indicating their potential to elicit xenogeneic immune reactions.

The identification of glycoproteins among the enriched proteins further emphasized their potential immunogenicity. Glycoproteins are known to play significant roles in immune recognition and response due to their glycan structures. Among the identified intact glycopeptides, 31 belonged to haptoglobin. Haptoglobin is a glycoprotein known for its immune-related functions, including binding free hemoglobin to prevent oxidative damage^43^. Previous studies have highlighted the influence of haptoglobin on transplant outcomes, demonstrating its role in activating innate immunity, which can lead to an enhancement of acute transplant rejection^44–46^. Despite these studies highlighting haptoglobin’s value in the field of organ transplantation, its potential to act as a xenoantigen has never been discovered or investigated. Our results suggest a new perspective by indicating that the enrichment of haptoglobin glycopeptides implies this protein may act as a xenoantigen in humans.

In conclusion, we have developed a novel proteomic approach for identifying xenoantigens. Using this approach, we identified 221 peptides corresponding to 153 proteins in the proteomic data, with a significant enrichment of plasma membrane and extracellular proteins, including 11 peptides unique to pig sequences and 122 intact glycopeptides corresponding to 16 glycoproteins in the glycoproteomic data, predominantly complex/hybrid glycoforms and Neu5Gc-containing glycans. These findings provide insights into discovering xenoantigens and warrant further investigation into identifying cytotoxic xenoantigens that lead to antibody-mediated rejection. Confirmation of cytotoxic xenoantigens could inform genetic engineering strategies and improve recipient candidate screening for xenotransplantation.

## ASSOCIATED CONTENT

Supplementary Figures: Figure S1. The efficiency of antibody recovery. The bar graph illustrated the efficiency of antibody recovery using varying volumes of pig kidney protein (PKP) linked beads and human pooled plasma. The x-axis represents different ratios of PKP beads to plasma tested: 100 μl and 200 μl of PKP beads with plasma volumes of 250 μl, 500 μl, and 1000 μl. The y-axis shows the amounts of recovered antibodies. Figure S2. The distribution of fold changes in protein- and peptide-level enrichments. A. Histograms and density plots compared the distribution of fold changes in protein-level enrichments. Red Bar Plots: Fold change distribution of the experimental group (PK, beads with specific antibodies to pig kidney proteins) compared to their own average values, showing the data distribution. Blue Bar Plots: Fold change distribution of the PK group compared to the control group (E, beads with non-specific antibodies), indicating the enrichment significance. Purple Dashed Lines: Intersection of the two fitting curves used as the cutoff. B. Histograms and density plots compared the distribution of fold changes in peptide-level enrichments. Red Bar Plots: Fold change distribution of the experimental group (PK, beads with specific antibodies to pig kidney proteins) compared to their own average values, showing the data distribution. Blue Bar Plots: Fold change distribution of the PK group compared to the control group (E, beads with non-specific antibodies), indicating the enrichment significance. Purple Dashed Lines: Intersection of the two fitting curves used as the cutoff.

Supplementary Tables: Table S1. The efficiency of antibody recovery. Table S2. The correlation results of protein and peptide-level enrichment. The data of the overlap of proteins enriched across three replicate samples (PK1, PK2, PK3) at the protein-level enrichment or peptide-level enrichment. The data of the overlap between proteins identified at the protein-level enrichment and proteins enriched at the peptide-level enrichment. Table S3. Intersections of protein and peptide-level enriched xenoantigens across cellular components and sequence alignment results. Table S4. Glycopeptide analysis results of the peptide-level enrichment.

## DATA ACCESSIBILITY

The mass spectrometry proteomics data have been deposited to the ProteomeXchange Consortium via the PRIDE^47^ partner repository with the data set identifier PXD071660. Reviewers can access the dataset by logging in to the PRIDE website using the following account details: Username: Password: P7it99Q0Ubb3

## ACKNOWLEDGMENT

This work was supported by the National Institutes of Health, National Cancer Institute, and the Clinical Proteomic Tumor Analysis Consortium (CPTAC, U24CA271079). This manuscript is the result of funding in whole or in part by the National Institutes of Health (NIH). It is subject to the NIH Public Access Policy. Through acceptance of this federal funding, NIH has been given a right to make this manuscript publicly available in PubMed Central upon the Official Date of Publication, as defined by NIH.

## For TOC Only

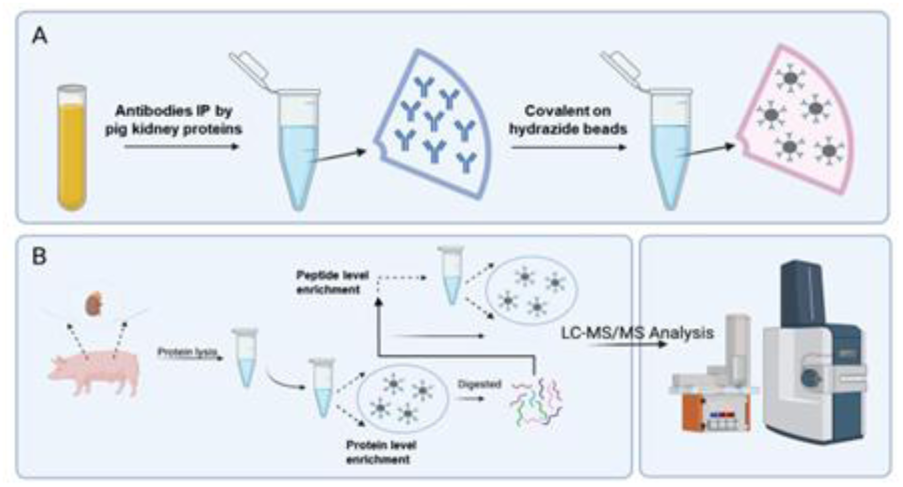

## References

(1) Vanderpool, H. Y. Xenotransplantation: Progress and Promise. BMJ 1999, 319 (7220), 1311.

(2) Yang, Y.-G.; Sykes, M. Xenotransplantation: Current Status and a Perspective on the Future. Nat. Rev. Immunol. 2007, 7 (7), 519–531. 10.1038/nri2099.

(3) Ekser, B.; Li, P.; Cooper, D. K. C. Xenotransplantation: Past, Present, and Future. Curr. Opin. Organ Transplant. 2017, 22 (6), 513. 10.1097/MOT.0000000000000463.

(4) Cascalho, M.; Platt, J. L. The Immunological Barrier to Xenotransplantation. Immunity 2001, 14 (4), 437–446. 10.1016/S1074-7613(01)00124-8.

(5) Vadori, M.; Cozzi, E. Current Challenges in Xenotransplantation. Curr. Opin. Organ Transplant. 2024, 29 (3), 205–211. 10.1097/MOT.0000000000001146.

(6) Galili, U. Interaction of the Natural Anti-Gal Antibody with α-Galactosyl Epitopes: A Major Obstacle for Xenotransplantation in Humans. Immunol. Today 1993, 14 (10), 480–482. 10.1016/0167-5699(93)90261-I.

(7) Byrne, G.; Ahmad-Villiers, S.; Du, Z.; McGregor, C. B4GALNT2 and Xenotransplantation: A Newly Appreciated Xenogeneic Antigen. Xenotransplantation 2018, 25 (5), e12394. 10.1111/xen.12394.

(8) Tector, A. J.; Mosser, M.; Tector, M.; Bach, J.-M. The Possible Role of Anti-Neu5Gc as an Obstacle in Xenotransplantation. Front. Immunol. 2020, 11. 10.3389/fimmu.2020.00622.

(9) Estrada, J. L.; Martens, G.; Li, P.; Adams, A.; Newell, K. A.; Ford, M. L.; Butler, J. R.; Sidner, R.; Tector, M.; Tector, J. Evaluation of Human and Non-Human Primate Antibody Binding to Pig Cells Lacking GGTA1/CMAH/β4GalNT2 Genes. Xenotransplantation 2015, 22 (3), 194–202. 10.1111/xen.12161.

(10) Cooper, D. K. C.; Hara, H.; Iwase, H.; Yamamoto, T.; Wang, Z.-Y.; Jagdale, A.; Bikhet, M. H.; Nguyen, H. Q.; Foote, J. B.; Paris, W. D.; Ayares, D.; Kumar, V.; Anderson, D. J.; Locke, J. E.; Eckhoff, D. E. Pig Kidney Xenotransplantation: Progress toward Clinical Trials. Clin. Transplant. 2021, 35 (1), e14139. 10.1111/ctr.14139.

(11) Reardon, S. First Pig-to-Human Heart Transplant: What Can Scientists Learn? Nature 2022, 601 (7893), 305–306. 10.1038/d41586-022-00111-9.

(12) Mallapaty, S.; Kozlov, M. First Pig Kidney Transplant in a Person: What It Means for the Future. Nature 2024, 628 (8006), 13–14. 10.1038/d41586-024-00879-y.

(13) Kozlov, M. Pig-Organ Transplants: What Three Human Recipients Have Taught Scientists. Nature 2024, 629 (8014), 980–981. 10.1038/d41586-024-01453-2.

(14) Montgomery, R. A.; Stern, J. M.; Lonze, B. E.; Tatapudi, V. S.; Mangiola, M.; Wu, M.; Weldon, E.; Lawson, N.; Deterville, C.; Dieter, R. A.; Sullivan, B.; Boulton, G.; Parent, B.; Piper, G.; Sommer, P.; Cawthon, S.; Duggan, E.; Ayares, D.; Dandro, A.; Fazio-Kroll, A.; Kokkinaki, M.; Burdorf, L.; Lorber, M.; Boeke, J. D.; Pass, H.; Keating, B.; Griesemer, A.; Ali, N. M.; Mehta, S. A.; Stewart, Z. A. Results of Two Cases of Pig-to-Human Kidney Xenotransplantation. N. Engl. J. Med. 2022, 386 (20), 1889–1898. 10.1056/NEJMoa2120238.

(15) Loupy, A.; Goutaudier, V.; Giarraputo, A.; Mezine, F.; Morgand, E.; Robin, B.; Khalil, K.; Mehta, S.; Keating, B.; Dandro, A.; Certain, A.; Tharaux, P.-L.; Narula, N.; Tissier, R.; Giraud, S.; Hauet, T.; Pass, H. I.; Sannier, A.; Wu, M.; Griesemer, A.; Ayares, D.; Tatapudi, V.; Stern, J.; Lefaucheur, C.; Bruneval, P.; Mangiola, M.; Montgomery, R. A. Immune Response after Pig-to-Human Kidney Xenotransplantation: A Multimodal Phenotyping Study. The Lancet 2023, 402 (10408), 1158–1169. 10.1016/S0140-6736(23)01349-1.

(16) Cooper, D. K. C.; Habibabady, Z.; Kinoshita, K.; Hara, H.; Pierson, R. N. The Respective Relevance of Sensitization to Alloantigens and Xenoantigens in Pig Organ Xenotransplantation. Hum. Immunol. 2023, 84 (1), 18–26. 10.1016/j.humimm.2022.06.003.

(17) Ladowski, J. M.; Hara, H.; Cooper, D. K. C. The Role of SLAs in Xenotransplantation. Transplantation 2021, 105 (2), 300–307. 10.1097/TP.0000000000003303.

(18) Lucander, A. C. K.; Nguyen, H.; Foote, J. B.; Cooper, D. K. C.; Hara, H. Immunological Selection and Monitoring of Patients Undergoing Pig Kidney Transplantation. Xenotransplantation 2021, 28 (4), e12686. 10.1111/xen.12686.

(19) Yamamoto, T.; Iwase, H.; Patel, D.; Jagdale, A.; Ayares, D.; Anderson, D.; Eckhoff, D. E.; Cooper, D. K. C.; Hara, H. Old World Monkeys Are Less than Ideal Transplantation Models for Testing Pig Organs Lacking Three Carbohydrate Antigens (Triple-Knockout). Sci. Rep. 2020, 10 (1), 9771. 10.1038/s41598-020-66311-3.

(20) Gao, B.; Long, C.; Lee, W.; Zhang, Z.; Gao, X.; Landsittel, D.; Ezzelarab, M.; Ayares, D.; Huang, Y.; Cooper, D. K. C.; Wang, Y.; Hara, H. Anti-Neu5Gc and Anti-Non-Neu5Gc Antibodies in Healthy Humans. PloS One 2017, 12 (7), e0180768. 10.1371/journal.pone.0180768.

(21) Zhang, M.; Ou, X.; Shi, H.; Huang, W.; Song, L.; Zhu, J.; Yu, R. Isolation, Structures and Biological Activities of Medicinal Glycoproteins from Natural Resources: A Review. Int. J. Biol. Macromol. 2023, 244, 125406. 10.1016/j.ijbiomac.2023.125406.

(22) Zhang, T.; Pierson, R. N.; Azimzadeh, A. M. Update on CD40 and CD154 Blockade in Transplant Models. Immunotherapy 2015, 7 (8), 899–911. 10.2217/IMT.15.54.

(23) Higginbotham, L.; Mathews, D.; Breeden, C. A.; Song, M.; Farris, A. B.; Larsen, C. P.; Ford, M. L.; Lutz, A. J.; Tector, M.; Newell, K. A.; Tector, A. J.; Adams, A. B. Pre-Transplant Antibody Screening and Anti-CD154 Costimulation Blockade Promote Long-Term Xenograft Survival in a Pig-to-Primate Kidney Transplant Model. Xenotransplantation 2015, 22 (3), 221–230. 10.1111/xen.12166.

(24) Aebersold, R.; Mann, M. Mass-Spectrometric Exploration of Proteome Structure and Function. Nature 2016, 537 (7620), 347–355. 10.1038/nature19949.

(25) Yates, J. R. Recent Technical Advances in Proteomics. F1000Research 2019, 8, F1000 Faculty Rev-351. 10.12688/f1000research.16987.1.

(26) Gillet, L. C.; Navarro, P.; Tate, S.; Röst, H.; Selevsek, N.; Reiter, L.; Bonner, R.; Aebersold, R. Targeted Data Extraction of the MS/MS Spectra Generated by Data-Independent Acquisition: A New Concept for Consistent and Accurate Proteome Analysis. Mol. Cell. Proteomics MCP 2012, 11 (6), O111.016717. 10.1074/mcp.O111.016717.

(27) Meier, F.; Brunner, A.-D.; Frank, M.; Ha, A.; Bludau, I.; Voytik, E.; Kaspar-Schoenefeld, S.; Lubeck, M.; Raether, O.; Bache, N.; Aebersold, R.; Collins, B. C.; Röst, H. L.; Mann, M. diaPASEF: Parallel Accumulation–Serial Fragmentation Combined with Data-Independent Acquisition. Nat. Methods 2020, 17 (12), 1229–1236. 10.1038/s41592-020-00998-0.

(28) Hu, Y.; Schnaubelt, M.; Chen, L.; Zhang, B.; Hoang, T.; Lih, T. M.; Zhang, Z.; Zhang, H. MS-PyCloud: A Cloud Computing-Based Pipeline for Proteomic and Glycoproteomic Data Analyses. Anal. Chem. 2024, 96 (25), 10145–10151. 10.1021/acs.analchem.3c01497.

(29) Sun, Z.; Lih, T. M.; Woo, J.; Jiao, L.; Hu, Y.; Wang, Y.; Liu, H.; Zhang, H. Improving Glycoproteomic Analysis Workflow by Systematic Evaluation of Glycopeptide Enrichment, Quantification, Mass Spectrometry Approach, and Data Analysis Strategies. Anal. Chem. 2024, 96 (52), 20481–20490. 10.1021/acs.analchem.4c04466.

(30) Berggård, T.; Linse, S.; James, P. Methods for the Detection and Analysis of Protein-Protein Interactions. Proteomics 2007, 7 (16), 2833–2842. 10.1002/pmic.200700131.

(31) Free, R. B.; Hazelwood, L. A.; Sibley, D. R. Identifying Novel Protein-Protein Interactions Using Co-Immunoprecipitation and Mass Spectroscopy. Curr. Protoc. Neurosci. Editor. Board Jacqueline N Crawley Al 2009, 0 5, Unit-5.28. 10.1002/0471142301.ns0528s46.

(32) Budayeva, H. G.; Cristea, I. M. A Mass Spectrometry View of Stable and Transient Protein Interactions. Adv. Exp. Med. Biol. 2014, 806, 263–282. 10.1007/978-3-319-06068-2_11.

(33) Immunoprecipitation: Techniques, Applications, and Future Challenges - Creative Proteomics. https://www.creative-proteomics.com/resource/immunoprecipitation-techniques-applications-future-challenges.htm (accessed 2025-01-02).

(34) Hughes, C. S.; Foehr, S.; Garfield, D. A.; Furlong, E. E.; Steinmetz, L. M.; Krijgsveld, J. Ultrasensitive Proteome Analysis Using Paramagnetic Bead Technology. Mol. Syst. Biol. 2014, 10 (10), 757. 10.15252/msb.20145625.

(35) Sinz, A. Cross-Linking/Mass Spectrometry for Studying Protein Structures and Protein–Protein Interactions: Where Are We Now and Where Should We Go from Here? Angew. Chem. Int. Ed. 2018, 57 (22), 6390–6396. 10.1002/anie.201709559.

(36) Sun, S.; Shah, P.; Eshghi, S. T.; Yang, W.; Trikannad, N.; Yang, S.; Chen, L.; Aiyetan, P.; Höti, N.; Zhang, Z.; Chan, D. W.; Zhang, H. Comprehensive Analysis of Protein Glycosylation by Solid-Phase Extraction of N-Linked Glycans and Glycosite-Containing Peptides. Nat. Biotechnol. 2016, 34 (1), 84–88. 10.1038/nbt.3403.

(37) Zhang, H.; Li, X.; Martin, D. B.; Aebersold, R. Identification and Quantification of N-Linked Glycoproteins Using Hydrazide Chemistry, Stable Isotope Labeling and Mass Spectrometry. Nat. Biotechnol. 2003, 21 (6), 660–666. 10.1038/nbt827.

(38) Bobbitt, J. M. Periodate Oxidation of Carbohydrates. In Advances in Carbohydrate Chemistry; Wolfrom, M. L., Tipson, R. S., Eds.; Academic Press, 1956; Vol. 11, pp 1–41. 10.1016/S0096-5332(08)60115-0.

(39) Altman, M. O.; Gagneux, P. Absence of Neu5Gc and Presence of Anti-Neu5Gc Antibodies in Humans—An Evolutionary Perspective. Front. Immunol. 2019, 10, 789. 10.3389/fimmu.2019.00789.

(40) Seo, N.; Ko, J.; Lee, D.; Jeong, H.; Oh, M. J.; Kim, U.; Lee, D. H.; Kim, J.; Choi, Y. J.; An, H. J. In-Depth Characterization of Non-Human Sialic Acid (Neu5Gc) in Human Serum Using Label-Free ZIC-HILIC/MRM-MS. Anal. Bioanal. Chem. 2021, 413 (20), 5227–5237. 10.1007/s00216-021-03495-1.

(41) Wylie, A. D.; Zandberg, W. F. Quantitation of Sialic Acids in Infant Formulas by Liquid Chromatography-Mass Spectrometry: An Assessment of Different Protein Sources and Discovery of New Analogues. J. Agric. Food Chem. 2018, 66 (30), 8114–8123. 10.1021/acs.jafc.8b01042.

(42) Ding, F.; Lin, Y.; Liu, G.; Liu, Y.; Gao, F.; Liu, Q.; Zhang, Z.; Weng, S. Immune Disguise: The Mechanisms of Neu5Gc Inducing Autoimmune and Transplant Rejection. Genes Immun. 2022, 23 (6), 175–182. 10.1038/s41435-022-00182-8.

(43) Andersen, C. B. F.; Stødkilde, K.; Sæderup, K. L.; Kuhlee, A.; Raunser, S.; Graversen, J. H.; Moestrup, S. K. Haptoglobin. Antioxid. Redox Signal. 2017, 26 (14), 814–831. 10.1089/ars.2016.6793.

(44) Shen, H.; Heuzey, E.; Mori, D. N.; Wong, C. K.; Colangelo, C. M.; Chung, L. M.; Bruce, C.; Slizovskiy, I. B.; Booth, C. J.; Kreisel, D.; Goldstein, D. R. Haptoglobin Enhances Cardiac Transplant Rejection. Circ. Res. 2015, 116 (10), 1670–1679. 10.1161/CIRCRESAHA.116.305406.

(45) Shen, H.; Song, Y.; Colangelo, C. M.; Wu, T.; Bruce, C.; Scabia, G.; Galan, A.; Maffei, M.; Goldstein, D. R. Haptoglobin Activates Innate Immunity to Enhance Acute Transplant Rejection in Mice. J. Clin. Invest. 2012, 122 (1), 383–387. 10.1172/JCI58344.

(46) Redmond, A. K.; Ohta, Y.; Criscitiello, M. F.; Macqueen, D. J.; Flajnik, M. F.; Dooley, H. Haptoglobin Is a Divergent MASP Family Member That Neofunctionalized To Recycle Hemoglobin via CD163 in Mammals. J. Immunol. 2018, 201 (8), 2483–2491. 10.4049/jimmunol.1800508.

(47) Perez-Riverol, Y.; Bai, J.; Bandla, C.; García-Seisdedos, D.; Hewapathirana, S.; Kamatchinathan, S.; Kundu, D. J.; Prakash, A.; Frericks-Zipper, A.; Eisenacher, M.; Walzer, M.; Wang, S.; Brazma, A.; Vizcaíno, J. A. The PRIDE Database Resources in 2022: A Hub for Mass Spectrometry-Based Proteomics Evidences. Nucleic Acids Res. 2022, 50 (D1), D543–D552. 10.1093/nar/gkab1038.

