## Supplemental Figures for "Proteomic Identification of Pig Xenoantigens for Clinical Xenotransplantation"

**Table of contents**

**Figure S1.** The efficiency of antibody recovery. ........................................................................S3
**Figure S2.** The distribution of fold changes in protein- and peptide-level enrichments.............S4

The efficiency of antibody recovery, related to Figure S1. (**Table S1**) (.xlsx)

The correlation results of protein and peptide-level enrichment. The data of the overlap of proteins enriched across three replicate samples (PK1, PK2, PK3) at the protein-level enrichment or peptide-level enrichment. The data of the overlap between proteins identified at the protein-level enrichment and proteins enriched at the peptide-level enrichment, related to Figure 2. (**Table S2**) (.xlsx)

Intersection analysis of protein and peptide-level enriched xenoantigens across cellular components and sequence alignment results, related to Figure 3. (**Table S3**) (.xlsx)

Glycopeptide analysis of the peptide-level enrichment, related to Figure 4. (**Table S4**) (.xlsx)

### Figure S1

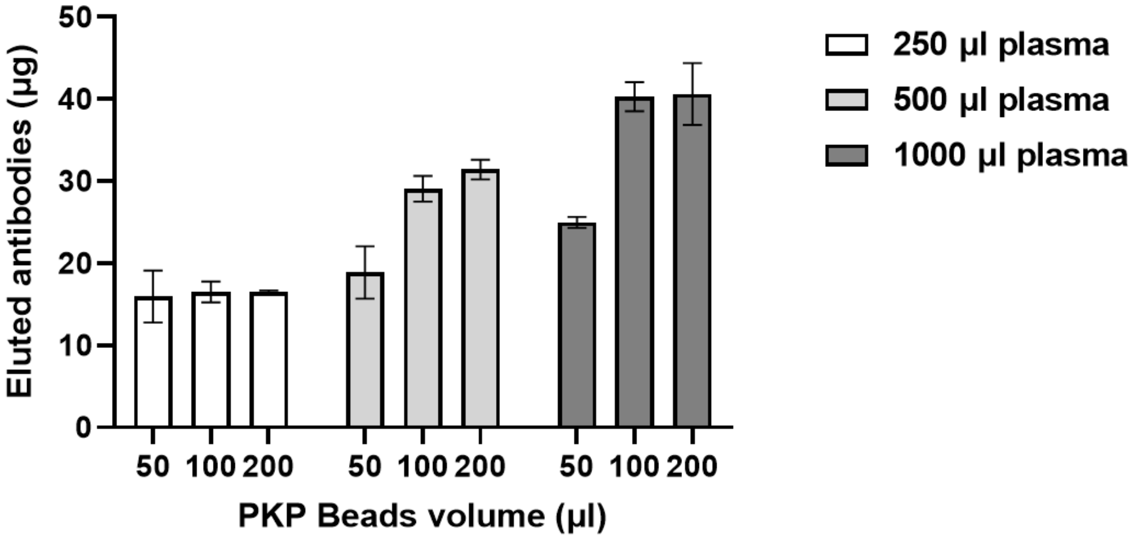


The bar graph illustrated the efficiency of antibody recovery using varying volumes of pig kidney protein (PKP) linked beads and human pooled plasma. The x-axis represents different ratios of PKP beads to plasma tested: 100 µl and 200 µl of PKP beads with plasma volumes of 250 µl, 500 µl, and 1000 µl. The y-axis shows the amounts of recovered antibodies.

### Figure S2

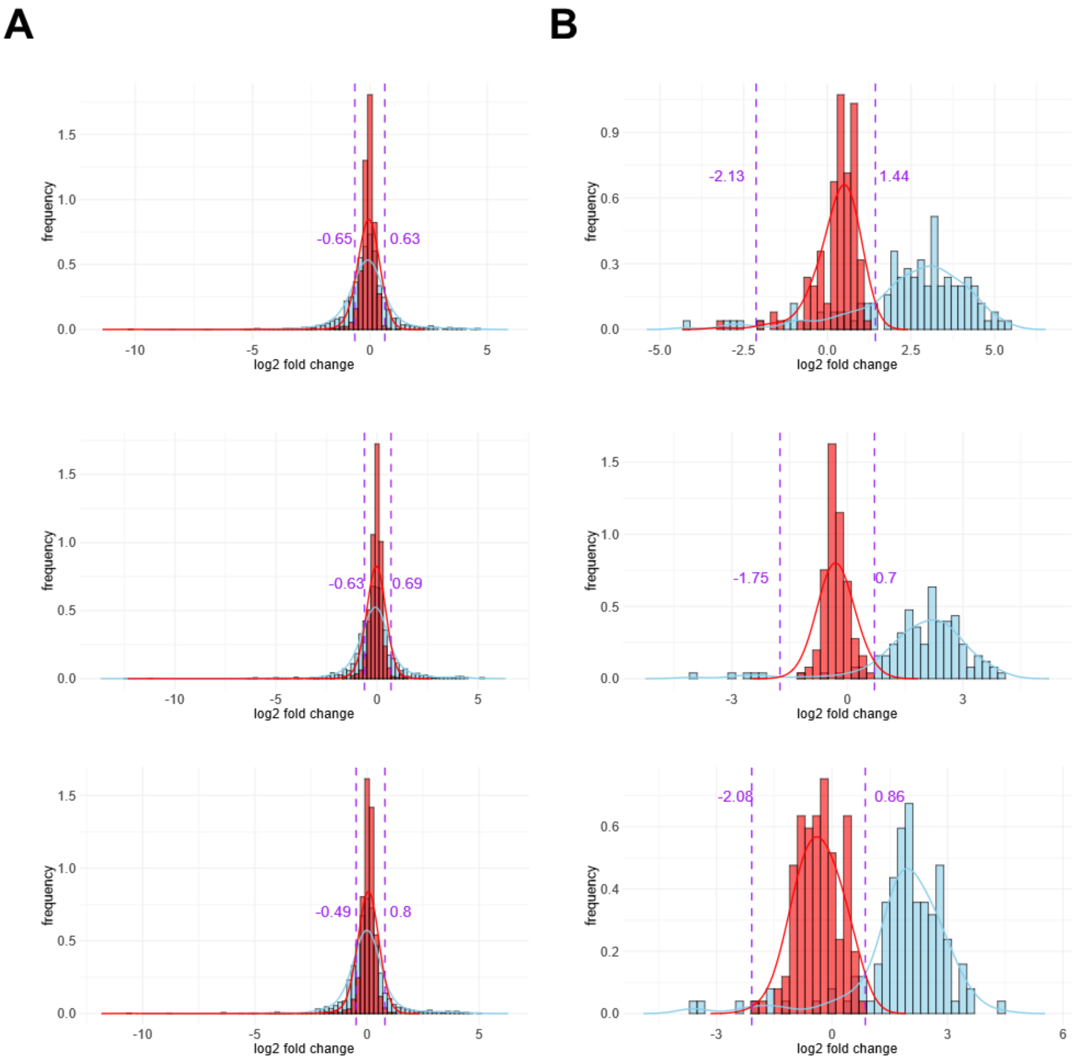


A. Histograms and density plots compared the distribution of fold changes in protein-level enrichments. Red Bar Plots: Fold change distribution of the experimental group (PK, beads with specific antibodies to pig kidney proteins) compared to their own average values, showing the data distribution. Blue Bar Plots: Fold change distribution of the PK group compared to the control group (E, beads with non-specific antibodies), indicating the enrichment significance. Purple Dashed Lines: Intersection of the two fitting curves used as the cutoff.

B. Histograms and density plots compared the distribution of fold changes in peptide-level enrichments. Red Bar Plots: Fold change distribution of the experimental group (PK, beads with specific antibodies to pig kidney proteins) compared to their own average values, showing the data distribution. Blue Bar Plots: Fold change distribution of the PK group compared to the control group (E, beads with non-specific antibodies), indicating the enrichment significance. Purple Dashed Lines: Intersection of the two fitting curves used as the cutoff.
